# Long Non-Coding RNAs in Response to Ebola Virus Vaccine-Induced Immunity

**DOI:** 10.1101/2025.08.19.671152

**Authors:** Izabela Mamede, Thomaz Luschër-Dias, Isabelle Franco Moscardini, Patrícia González, Barbara Marinho, Fernando Marcon, Thiago Dominguez Crespo Hirata, VSV-EBOPLUS Consortium, Michael Eichberg, Donata Medaglini, Ali M Harandi, Claire-Anne Siegrist, Tom H M Ottenhoff, Francesco Santoro, André Gonçalves, Rafael Polidoro, Glória R. Franco, Paulo P. Amaral, Helder Nakaya

## Abstract

Long noncoding RNAs (lncRNAs) have emerged as critical regulators of gene expression, yet their role in shaping human responses to vaccination remains largely uncharacterized. Here, we analyzed RNA-sequencing data from three independent human cohorts vaccinated with the rVSVΔG-ZEBOV-GP Ebola vaccine to profile lncRNA expression dynamics. Using differential expression analysis and correlation meta-analysis across cohorts, we identified an expression signature with several lncRNAs, including *LEF1-AS1* and *DOCK8-AS1*, that exhibit conserved transcriptional activation following vaccination. Correlation of lncRNA expression with gene targets and IgG titers revealed putative roles for lncRNAs in regulating and/or participate in both innate immune responses and adaptive antibody production. Functional enrichment of lncRNA co-expressed protein-coding genes highlighted involvement in T-cell differentiation, interferon signaling, and leukocyte activation. Integrating global run-on sequencing data and comparative transcriptomic analysis across other vaccine studies suggests that *LEF1-AS1* modulation is distinctively associated with Ebola vaccination. Our findings demonstrate that lncRNAs are potential integral components of the human vaccine response and provide a foundation for future mechanistic studies targeting noncoding RNA regulation of immunity

**Significance:** Ebola virus remains a significant global health threat due to its high mortality rate and potential for widespread outbreaks, underscoring the urgent need for effective and durable vaccines to control future epidemics. Understanding the transcriptional mechanisms underlying immune responses to vaccination is important to improving vaccine design and efficacy. While protein-coding genes have been extensively studied, the role of long noncoding RNAs (lncRNAs) in vaccine-induced immunity remains poorly understood. Here, we characterize the dynamics of lncRNA expression following administration of the rVSVΔG-ZEBOV-GP Ebola vaccine across multiple human cohorts and identify conserved lncRNA signatures associated with both innate and adaptive immunity.

**Graphical abstract:** 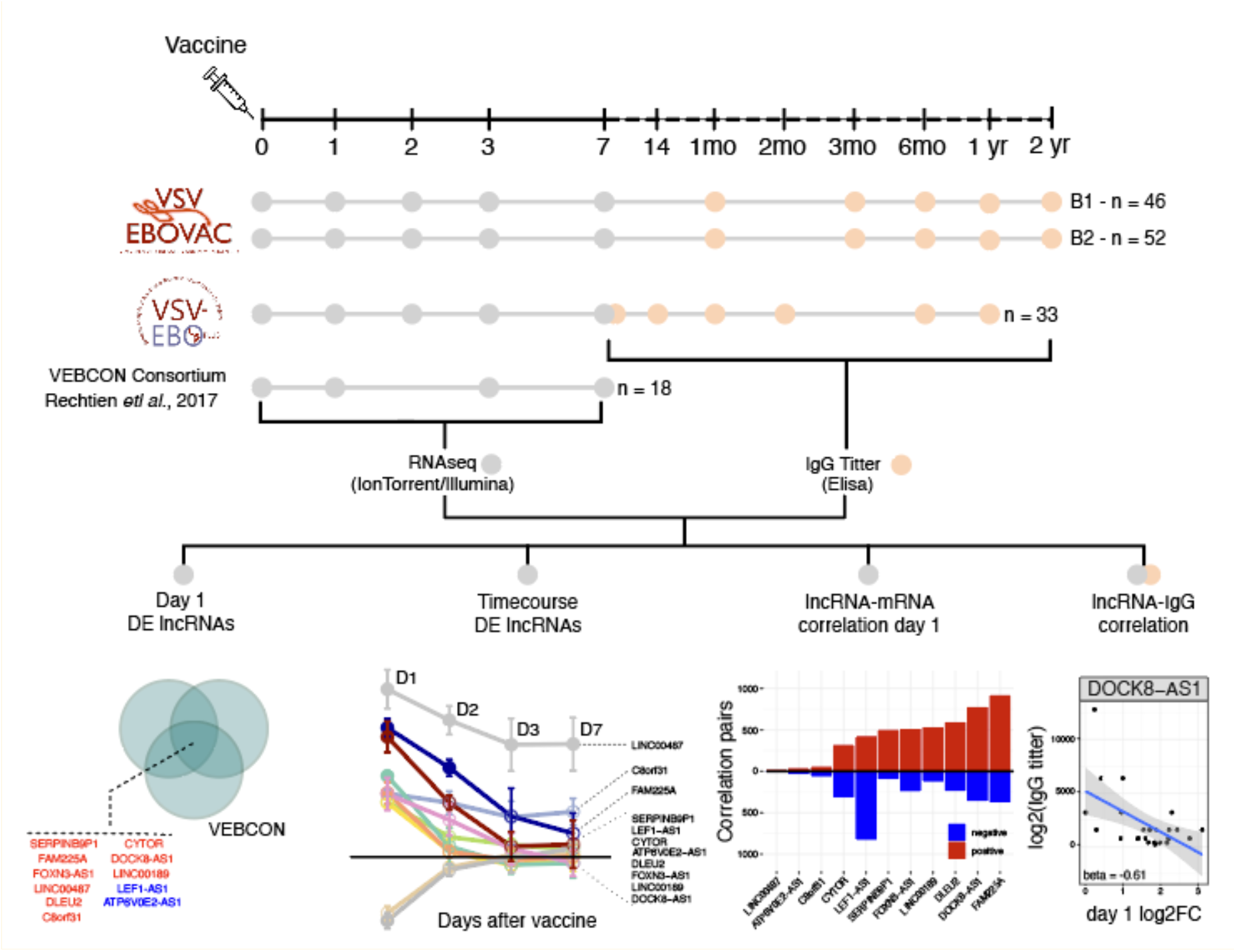

## Introduction

The Ebola virus disease (EVD) caused by *<u>Orthoebolavirus zairense</u>* is a severe hemorrhagic fever with a fatality rate ranging from 50% to 90%. The disease is highly contagious, and from 2014 to 2016, an Ebola outbreak in West Africa killed over 11,000 people, while between 2018 and 2020, the outbreak in the Democratic Republic of Congo (DRC) resulted in 3,323 cases with 2,299 deaths. In the aftermath of these outbreaks, coordinated efforts between research consortia, governments, global public health entities, and the private sector resulted in the approval of two treatments by the U.S. Food and Drug Administration (FDA) for EVD caused by Orthoebolavirus zairense; Inmazeb®, a combination of three monoclonal antibodies, and Ebanga®, a single monoclonal antibody (Food and Drug Administration [FDA], 2024); as well as the development of an efficacious and safe vaccine, the VSVΔG-ZEBOV-GP. This recombinant Vesicular Stomatitis Virus (VSV) expresses the Zaire Ebola virus (ZEBOV) glycoprotein (GP), and its efficacy was evaluated during the 2014–2016 outbreak in Guinea. The rVSV-ZEBOV is the first Ebola vaccine that is pre-approved to be used in response to Ebola outbreaks, and has been recently used, showing an 84% effectiveness (Meakin et al., 2024).

Systems vaccinology literature demonstrates the relevance of high-throughput techniques correlating gene expression before and after vaccination with higher quality and persistence of antibodies, guiding improvements in vaccine design and delivery(Vianello et al., 2022). VSV-EBOVAC Consortium and VSV EBOPLUS consortium sought to identify important correlates of protection during vaccination using VSVΔG-ZEBOV-GP in adult cohorts in Europe, Africa, and North America using transcriptomics (.They analyzed peripheral blood changes induced by the VSVΔG-ZEBOV-GP vaccine, which revealed that the peak of perturbation occurs one day after administration of the vaccine (Vianello et al., 2022). The study showed early immune response activation with increased type type I and II interferon response-related genes and myeloid cell-associated markers. Whereas, genes of circulating effector cells, such as cytotoxicity-associated genes, were downregulated in the peripheral blood cells of vaccinated individuals (Vianello et al., 2022). However, these analyses have primarily focused on protein-coding genes, leaving the role of non-coding RNA largely unexplored.

LncRNAs are long (>200 nucleotides) non-coding regulatory RNAs able to subtly regulate gene expression through RNA-RNA, RNA-DNA, and RNA-protein interactions(Mattick et al., 2023a). Our group and others have previously demonstrated the relevance of lncRNAs in immune modulation, including cytokine production (Obaid et al., 2018)myeloid cell activation (Wang et al., 2023) and T and B cell differentiation(Wang et al., 2022a). Several lncRNAs have been associated with immune responses to viral infections such as influenza, dengue, and yellow fever, and some are even being investigated as therapeutic targets in cancer and inflammatory diseases (Lüscher-Dias et al., 2021) Despite its potential in generating potent immunological memory (Chen et al., 2022; Wang et al., 2022a) the involvement of lncRNAs in the immune response to the Ebola vaccine has not been characterized. Investigating the LncRNAs changes in the vaccinated cohorts may reveal additional immune response mechanisms associated with improved outcomes following VSVΔG-ZEBOV-GP vaccination.

Here, we analyzed early lncRNA response upon Ebola vaccination and correlated their expression to IgG titer quantification obtained from three independent cohorts of volunteers vaccinated with the VSVΔG-ZEBOV-GP. The goal was to detect differentially expressed (DE) lncRNAs that could be mediating the long-term humoral protection conferred by the vaccine. In all cohorts, we detected 11 lncRNAs that were consistently differentially expressed on day 1 following the vaccination. Our analysis revealed three lncRNAs - *FAM225A, DOCK8-AS1*, and *LEF1-AS1* - with a robust correlation with IgG titers at several time points after the vaccine. Moreover, pairing affected pathways with differentially expressed lncRNAs suggested a potential regulatory role over their associated genes, which are involved in both innate and adaptive immunity.

## Results

### rVSV-ZEBOV vaccination induces consistent day 1 lncRNA expression across cohorts

We integrated three distinct cohorts of vaccinated volunteers in our study: VSV Ebovac (Geneva, n=98, two batches), VSV EBOPLUS (USA, n = 33), and VEBCON (Germany, n = 18) (Fig. 1) (Santoro Vaccines, Huttner Clin, Fischer Vaccine). Volunteers from all cohorts received one dose of the rVSV-ZEBOV vaccine on day zero and had their blood collected just before receiving the vaccine (day 0) and on subsequent days (1, 2, 3, and 7) for RNA sequencing analysis. Volunteers returned at several time points during the 2 years following the vaccination to provide blood for IgG titer quantification using ELISA. RNAseq and IgG titer results were then analyzed to obtain the consistently differentially expressed lncRNAs on day 1 and time course differential expression analysis of those lncRNAs, meta-analysis of the correlation between lncRNA and mRNA expression, and meta-analysis of the correlation between lncRNA expression and IgG titers.

**Figure 1:**
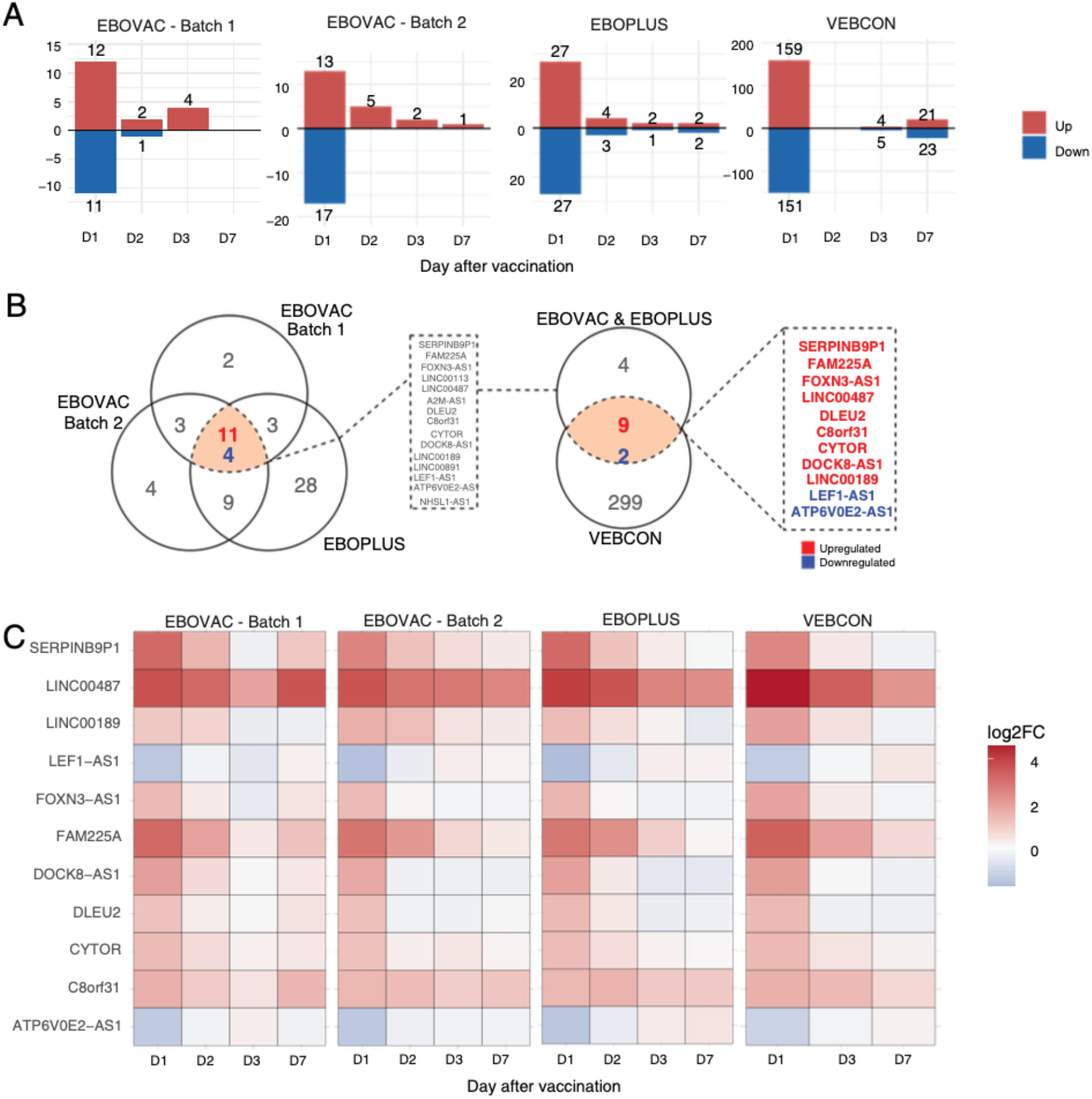
Selected lncRNA expression is consistent across time points and cohorts following rVSVΔG-ZEBOV-GP vaccination. **A**. Number of lncRNAs differentially expressed on days 1, 2, 3, and 7 after vaccination in the three different cohorts. **B**. Venn diagram representation of eleven lncRNAs consistently differentially expressed in all three vaccine cohorts on day 1 after vaccination. Blue: downregulated lncRNAs; red: upregulated lncRNAs. **C**. Heatmap analysis showing time course of the log_2_fold change of the 11 common day 1 DE lncRNAs shared between all cohorts.

In all cohorts, the peak of gene expression perturbation following vaccination occurred on day 1 **(Fig. 1A)**. On this day, the rVSV–ZEBOV vaccine elicited 12 up- and 11 downregulated lncRNAs for batch 1 EBOVAC volunteers, 13 up- and 17 downregulated lncRNAs for batch 2 EBOVAC volunteers, 27 up- and 27 downregulated lncRNAs for EBOPLUS volunteers, and 159 up- and 151 downregulated lncRNAs for VEBCON volunteers **(Fig. 1A)**. The number of differentially expressed lncRNAs in the Vebcon cohort was consistently higher than in the Ebovac and EBOPLUS cohorts, which can be attributed to the sequencing technology used for that cohort (Illumina) compared to that used in the other cohorts (IonTorrent). Since the vaccine transcriptomic perturbation was consistently higher on day 1 than that on days 2, 3, and 7 after the vaccine in all cohorts **(Fig. 1A)**, we decided to detect the lncRNAs that were consistently DE in all groups of volunteers on the first day post-vaccination **(Fig. 1B)**. Altogether, 15 lncRNAs (11 up- and 4 downregulated) were differentially expressed in common between the EBOVAC and EBOPLUS (batch 1 and batch 2) cohorts, 11 of which were also differentially expressed in the VEBCON cohort **(Fig. 1B, S1A)**. From these 11 consensus DE lncRNAs between the three cohorts, 9 were upregulated (*SERPINB9P1, FAM225A, FOXN3-AS1, LINC00487, DLEU2, C8orf31, CYTOR, DOCK8-AS1*, and *LINC00189*) and 2 were downregulated (*LEF1-AS1* and *ATP6V0E2-AS1*) **(Fig. 1B)**. The common lncRNAs that were mostly upregulated on day 1 were *LINC00487* and *FAM225A*, and the most downregulated one was *LEF1-AS1* **(Fig. 1C)**. The majority of the shared vaccine-induced lncRNAs returned to baseline levels on days 2, 3, and 7, except for *LINC00487*, which remained upregulated in all cohorts until day 7 **(Fig. 1C)**. On the three groups of volunteers which were analyzed on day 2, *FAM225A* was also upregulated, while this lncRNA was still upregulated on day 3 in the VEBCON patient cohort **(Fig. 1C, Fig S1B)**, but it also returned to baseline on day 7 post-vaccination.

**Figure S1:**
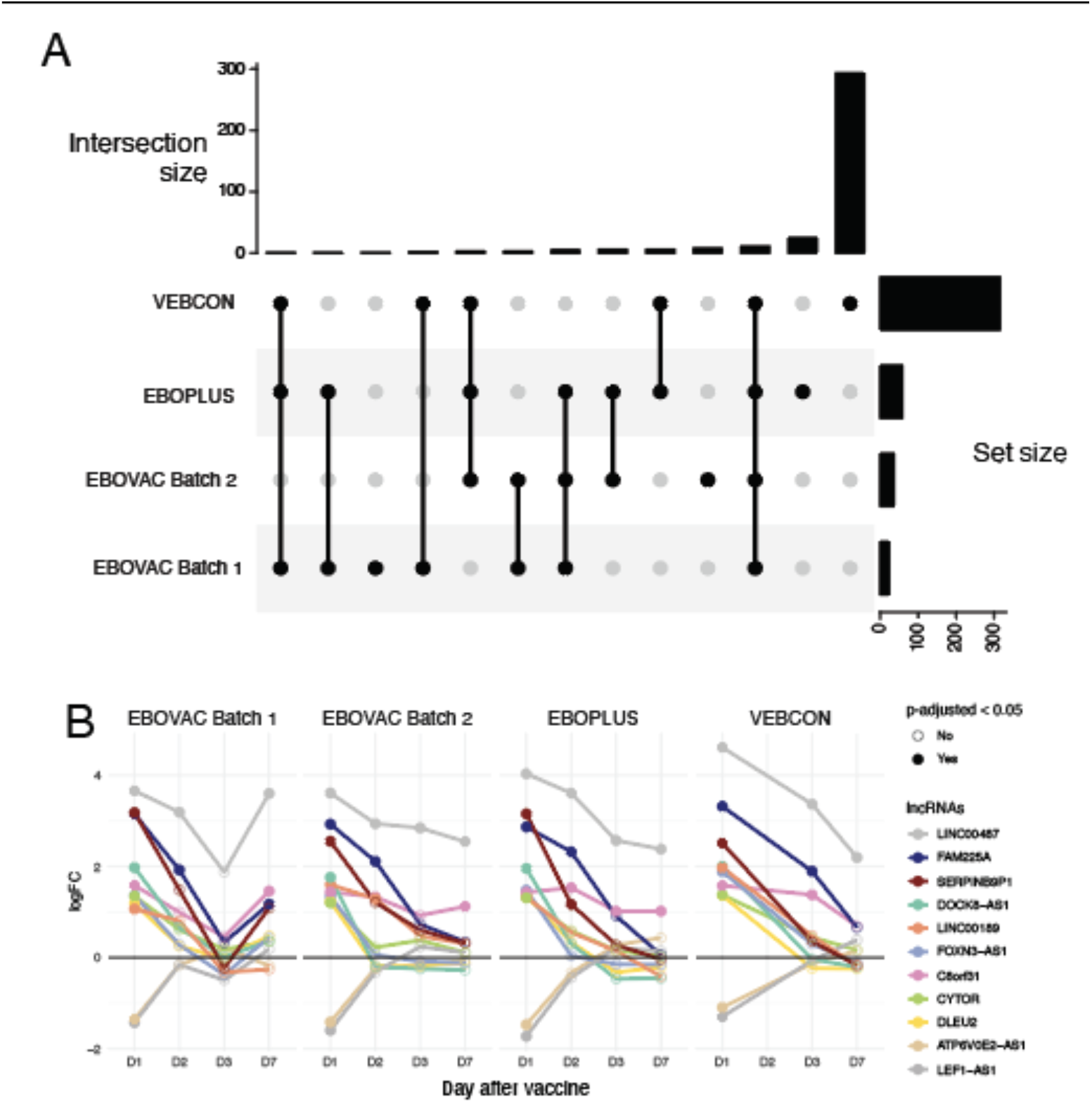
**A**. Upsetplot showing the intersection of differentially expressed lncRNAs across cohorts. **B**. Line graph of log FoldChange values switch of each selected lncRNA in all the 4 cohorts.

### Vaccine-induced lncRNAs show correlated expression with genes of key immune pathways and cells

lncRNAs can directly regulate the expression of protein-coding genes through lncRNA-RNA, lncRNA-DNA, or lncRNA-protein interactions (Mattick et al., 2023b). This regulation is particularly relevant in other immune-related phenotypes like viral and bacterial infections, where specific lncRNAs interact with viral components or immune regulators to modulate interferon responses, inflammation, antigen presentation, and even viral replication. To explore whether consensus day 1 VSV-ZEBOV-affected lncRNAs could be influencing the expression of other immune-related genes, we performed a correlation analysis between the expression of the 11 lncRNAs at day 1 and the expression of all other genes in the genome in each cohort and integrated those results using a meta-analysis approach.

The lncRNAs that showed the highest number of meta-correlated mRNAs on day 1 in all three cohorts were *DOCK8-AS1* (1,123 correlated mRNAs), *LEF1-AS1* (1,243 correlated mRNAs), and FAM225A (1,284 correlated mRNAs) **(Fig. 2A)**. *DOCK8-AS1* is encoded on chromosome 9 and is located on the opposite strand of the *DOCK8* (Dedicator of Cytokinesis 8) gene promoter, while *LEF1-AS1* is an anti-sense lncRNA located in chromosome 4 opposite to the protein-coding gene *LEF1* (Lymphoid Enhancer-Binding Factor 1). *FAM225A* is transcribed form the sense strand of chromosome 9 and its transcription have not been found in other mammalians.

**Figure 02:**
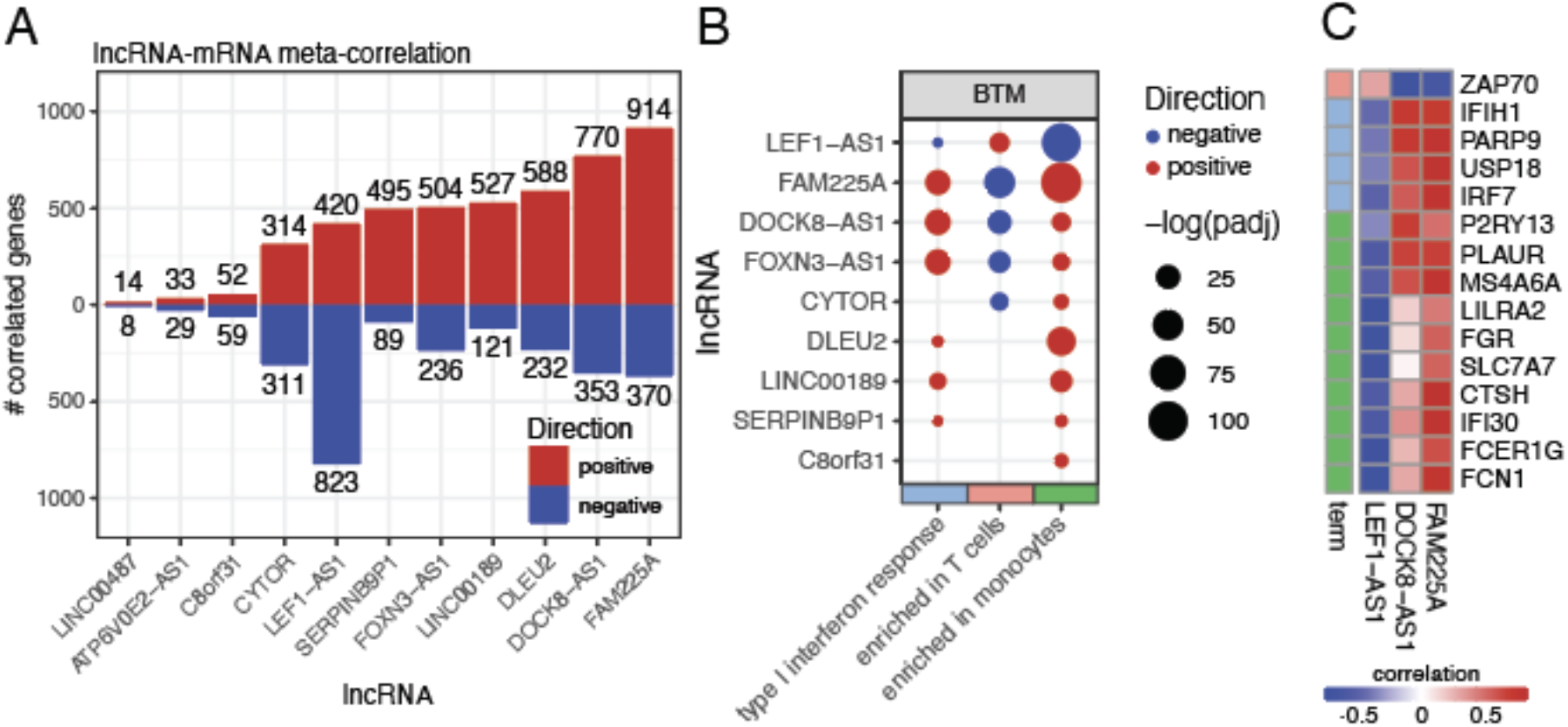
Day 1 vaccine-induced lncRNAs have correlated expression with key immune genes, pathways, and cells across cohorts. **A**. Number of meta-analysis significant correlation pairs between day 1 vaccine-induced lncRNAs and mRNAs. Blue: negative correlation pairs; red: positive correlation pairs. **B**. Overrepresentation analysis of blood transcriptional modules enriched among mRNAs positively (red) and negatively (blue) correlated with vaccine-induced lncRNAs. **C**. Top positive (red) and negative (blue) correlated mRNAs with lncRNAs FAM225A, LEF1-AS1, and DOCK8-AS1.

The mRNAs that positively correlated with *FAM225A* and *DOCK8-AS1* were functionally enriched for interferon response and monocytes, while the genes negatively correlated with these lncRNAs were enriched for genes commonly expressed by T cells **(Fig. 2B)**. Monocytes and interferon/innate response are the main upregulated pathways, while T cells are the main downregulated ones after rVSV-ZEBOV vaccination. Conversely, genes negatively correlated with LEF1-AS1 were enriched for interferon signaling and monocytes, and the genes positively correlated with LEF1-AS1 were enriched for T cell expression **(Fig 2B)**. Among the genes that showed a strong (r > 0.7) positive correlation with *FAM225* are *IRF7* **(Fig. 2C)**, a key mediator of the interferon signaling pathway, *IFIH1* (**Fig. 2C)**, the gene that codes for the viral sensing protein MDA5, and *USP18* **(Fig. 2C)**, a gene whose expression in macrophages has been shown to be essential for the effectiveness of the VSV-EBOV vaccine Friedrich Vaccines. *DOCK8-AS1* also presented a highly correlated expression with *IFIH1* and *P2RY13* **(Fig. 2C)**, an IFN-stimulated gene which protects against viral infections in mice(Zhang et al., 2019). Both *FAM225A* and *DOCK8-AS1* presented a negative correlation with *ZAP70* **(Fig. 2C)**, which codes for a membrane protein that participates in T-cell receptor (TCR)-mediated antigen-activation. LEF1-AS1, on the other hand, had a strong negative correlation with *FCN1* **(Fig. 2C)**, a gene coding for a collagen-like protein expressed in leukocytes, and *FCER1G* **(Fig. 2C)**, which codes for an Fc receptor of IgE.

### Vaccine-induced lncRNAs show correlated expression with IgG titers on several days after vaccination

Next, using the VSV-EBOVAC and VSV-EBOPLUS cohorts, we performed a meta-analysis of the correlation between the expression of the 11 VSV-ZEBOV-perturbed lncRNAs with circulating IgG titers in several time points after the vaccine. This correlation analysis was performed to identify potential regulatory roles of lncRNAs in shaping the humoral immune response elicited by the rVSV-ZEBOV vaccine. lncRNAs that exhibit significant correlation with IgG titers across multiple time points may contribute to B cell activation, differentiation, or antibody class switching essential to the vaccine response. Patient-level IgG titer data were not publicly available for the VEBCON cohort.

*FAM225A* expression on day 1 after the vaccine showed a positive correlation with antibody titers 14 **(Fig. 3A, B)** and 56 days post-vaccination **(Fig. 3A)**. Day 1 *DOCK8-AS1* levels were negatively correlated with IgG titers on days 28 **(Fig. 3C)**, 84, 168, and 365 **(Fig. 3A)**. *LEF1-AS1* was negatively correlated with IgG titers only on day 360 (1 year) **(Fig. 3D and 3A)**. *CYTOR* presented a negative correlation with antibody levels on days 28 and 168 (6 months) **(Fig. 3A)**, and *C8orf31* was positively correlated with IgG titers on days 14 and 56 (**Fig. 3A)**. IgG titers over time for lncRNAs *DOCK8-AS1* and *FAM225A* also showed significance and high correlation scores on other days without the meta-correlation **(Figure S2)**.

**Figure 3:**
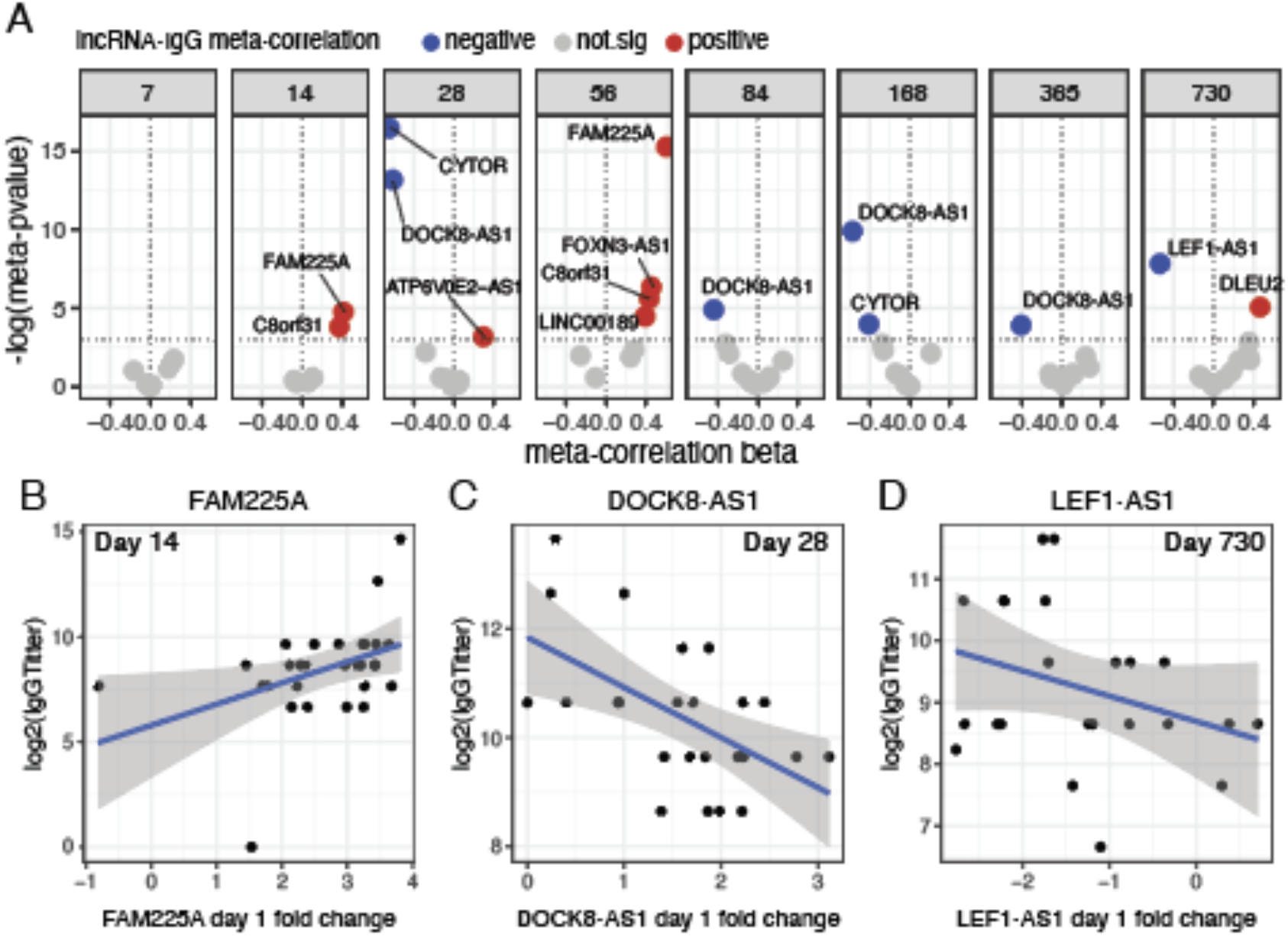
LncRNAs are correlated to IgG titer presence in volunteers after rVSVΔG-ZEBOV-GP vaccination. **A**. Significant meta-analysis correlations between day 1 vaccine-induced lncRNAs and IgG titer from 7 days to 2 years after vaccination. Meta-correlation p-values < 0.05 were considered significant. Individualized patient baseline normalized (log_2_-scaled D1-D0) expression of *FAM225A*; **B**., *DOCK8-AS1*; **C**., and *LEF1-AS1*; **D**. versus day 14, 28, and 730 IgG titers (log_2_ scale), respectively.

**Figure S2:**
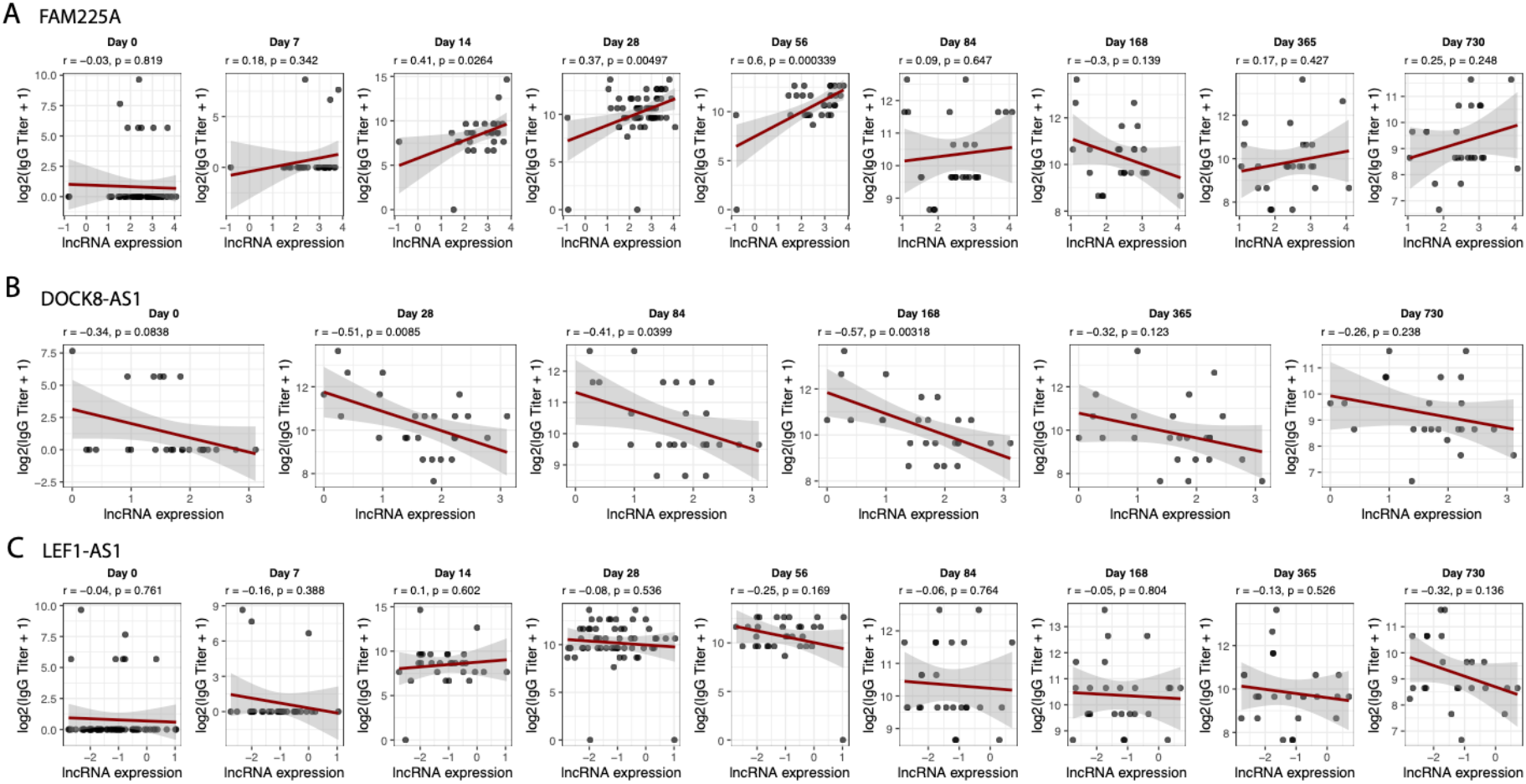
(A–C) Expression of three day 1–induced lncRNAs—*FAM225A* (A), *DOCK8-AS1* (B), and *LEF1-AS1* (C)—correlates with circulating anti-Ebola GP IgG titers across multiple time points after vaccination. Each subplot shows individualized volunteer expression (log2) versus IgG titer (log_2_[IgG + 1]) measured at the indicated day post-vaccination. Pearson correlation coefficients (r) and p-values are shown in each panel.

### *DOCK-AS1* can act as a promoter-antisense (PAS) RNA of the mRNA *DOCK8*

DOCK8 was one of the genes that presented the highest correlation with the lncRNA *DOCK8-AS1* in all days and cohorts **(Fig. 4A)**, being stronger on days 1 and 7 after the vaccination. This correlation was detected even on day 0, before vaccination **(Fig. 4A)**, which demonstrates that *DOCK8-AS1* and *DOCK8* likely have an intrinsically associated pattern of expression even in the absence of salient stimuli. Recently, evidence from Global run-on sequencing (GROseq) and CRISPR assays demonstrated that expression and shape of promoter antisense lncRNAs (PAS RNAs) are essential for the expression of their neighboring sense-encoded genes(Yang et al., 2021). Therefore, we hypothesized that *DOCK8-AS1* might be playing the role of a PAS RNA to induce the expression of *DOCK8* in vaccinated volunteers, leading to the correct migration of T_fh_ cells in response to the vaccination. We analyzed a publicly available GROseq dataset of human lymphoblasts infected with Sendai virus probed at several time points from 0 to 72h after infection (PRJEB22484) (Bitzer et al., 1999) **(Fig. 4B)**. One day after the infection, we detected an accumulation of reads across the entire DOCK8 gene on the sense strand **(Fig. 4B)**, but also a peak of reads mapping to the *DOCK8-AS1* gene on the antisense strand **(Fig. 4B)**. The GROseq assay allows for detection of co-occurring *de novo* nuclear RNA expression. This result implies that *DOCK8-AS1* and *DOCK8* are being co-expressed at the same time in the nucleus of lymphoblasts infected with Sendai virus. This pattern was detected across the other timepoints **(Fig. 4C)**, with evidence that Sendai virus infection is also capable of inducing increased expression of *DOCK8-AS1* and *DOCK8* 24 h after infection **(Fig. 4C)**, similarly to what we observed here for the VSV-ZEBOV Ebola vaccine in the blood.

**Figure 04:**
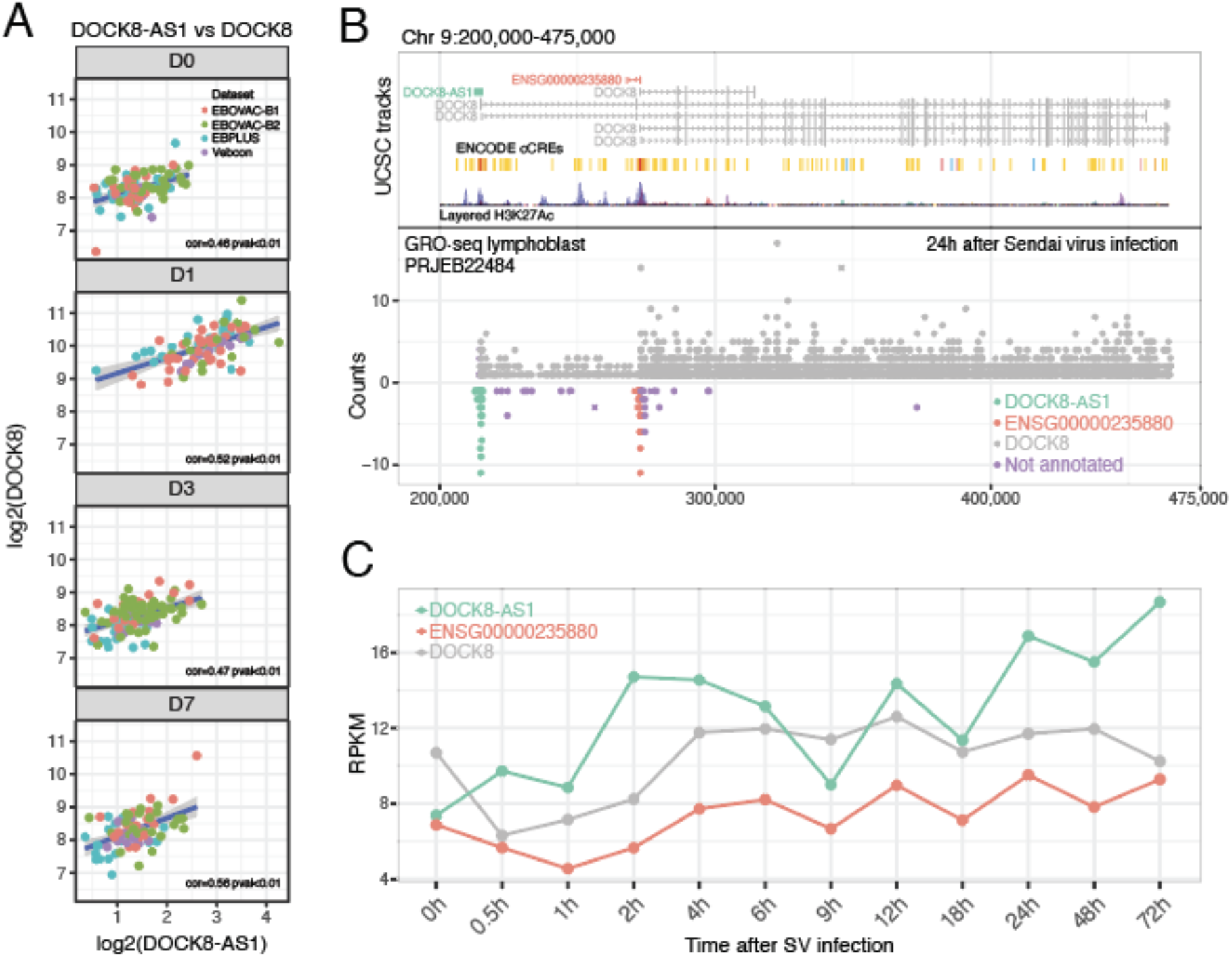
DOCK-AS1 is a putative promoter-antisense (PAS) RNA of the protein-coding gene DOCK8. **A**. *DOCK8-AS1* is positively correlated with *DOCK8* in all cohorts and days following rVSVΔG-ZEBOV-GP vaccination. **B**. Global Run-on sequencing (GRO-seq) analysis of lymphoblasts infected with Sendai virus after 24h shows concomitant peaks of expression of *DOCK8-AS1* (negative strand) and *DOCK8* (positive strand) around the *DOCK8* gene promoter region. **C**. *DOCK8* and *DOCK8-AS1* follow a similar pattern of co-expression through time after Sendai virus infection. *ENSG00000235880*, another lncRNA, follows a similar pattern.

Curiously, we also detected a peak of antisense reads mapping to the *ENSG00000235880* gene locus, in a region inside *DOCK8* with H3K27Ac epigenetic marks correlated with promoter activity **(Fig. 4B)**. Indeed, this position within the *DOCK8* gene coincides with the starting position of three shorter, alternative first exon isoforms of *DOCK8* **(Fig. 4C)**. Our analysis suggests that both *DOCK8-AS1* and *ENSG00000235880* might be acting as PAS RNAs within the canonical and alternative promoters of *DOCK8*, respectively.

### LEF1-AS1 controls the expression of mRNA LEF1 and is downregulated upon EBOLA vaccination

*LEF1-AS1* is an anti-sense lncRNA transcribed in chromosome 4 opposite to the protein-coding gene *LEF1* (Lymphoid Enhancer-Binding Factor 1). LEF1 translated protein itself is a positive regulator in lymphocyte development and activation(Reya et al., 2000), influencing the transcriptional programs that guide immune cell fate. Following rVSV-ZEBOV vaccination, *LEF1-AS1* expression sharply decreased on day 1, returning to baseline levels on days 2, 3, and 4. *LEF1* exhibited a matching expression profile, and its levels were strongly positively correlated with *LEF1-AS1* **(Fig. 5A, B)**. *LEF1-AS1* is transcribed from a region upstream of *LEF1* marked by H3K27Ac histone modifications and other promoter/enhancer marks **(Fig. 5B, C)**, suggesting a potential promoter-antisense (PAS) RNA regulatory role. Here we present evidence for *LEF1-AS1* ability to regulate the expression of *LEF1* (Qi et al., 2022). To assess whether this transcriptional regulation pattern was unique to the Ebola vaccine, we compared *LEF1-AS1* dynamics with transcriptomic data from other vaccine trials, including influenza and dengue. When looking at volunteers blood expression 3 days after Influenza vaccination in two separate cohorts (Moehling et al., 2020) or two days after dengue vaccination (Kim et al., 2022b), the same reduction pattern is not observed (**Figure 5D**). The influenza and dengue vaccines are based on inactivated and live attenuated viruses, respectively, suggesting that platform differences may underlie the distinct transcriptional responses elicited by the live replicating viral vector VSVΔG-ZEBOV-GP. Moreover, there was no dataset from Day 1 after vaccination transcriptomics available so a direct comparison with the day 1 potent immunogenic profile of the rVSV-ZEBOV vaccine was not possible. The coordinated downregulation of both *LEF1-AS1* and *LEF1* immediately after Ebola vaccination may reflect a transient suppression of Wnt signaling components to favor early innate immune responses or modulate antigen-specific T cell activation.

**Figure 5:**
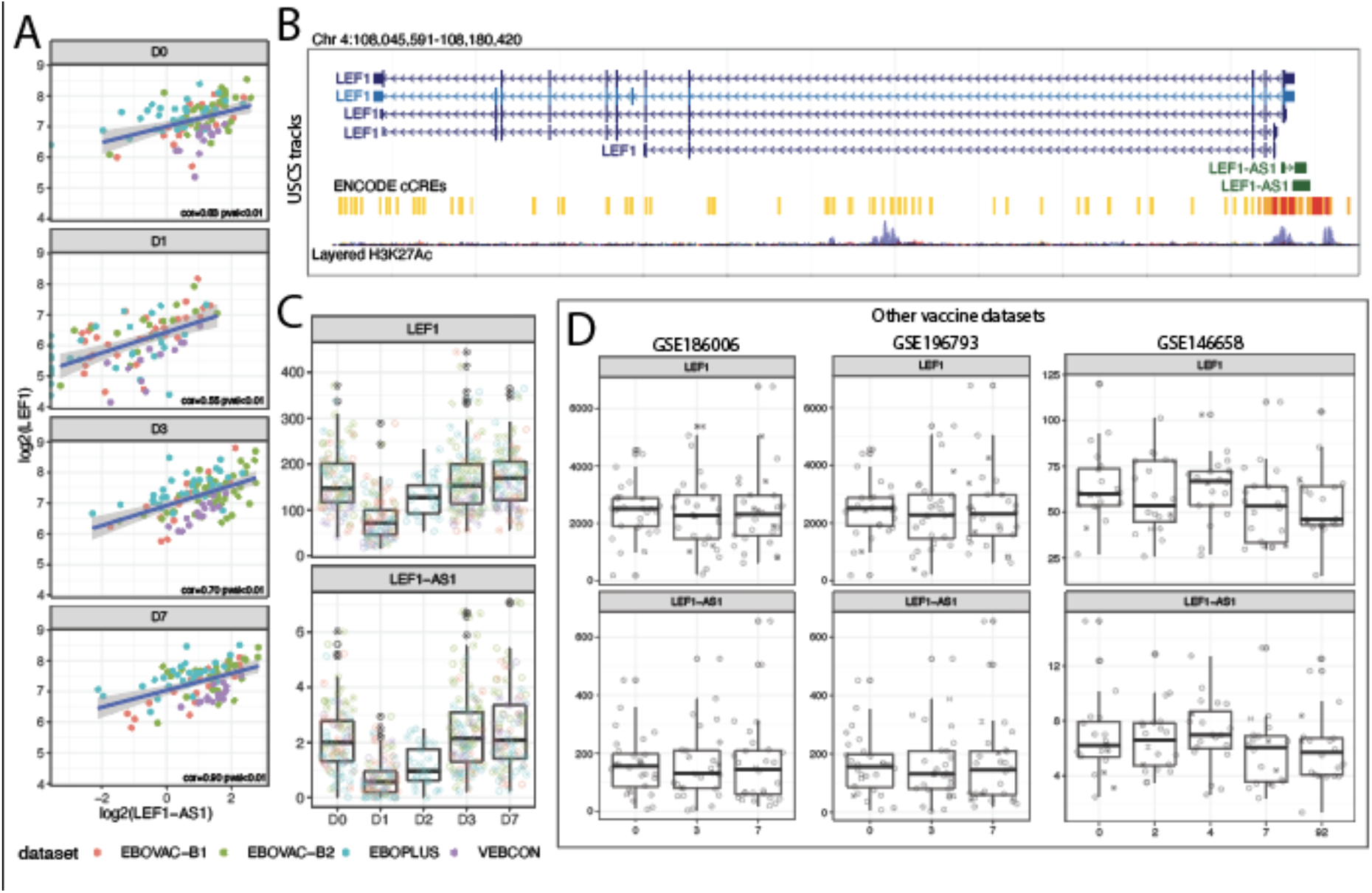
LEF1-AS1 is an anti-sense lncRNA that regulates LEF1 expression prominently on the first day after vaccination. **A**. Log2-transformed expression levels of *LEF1-AS1* and *LEF1* across days 0 to 7 post-vaccination in the four cohorts (red for EBOVAC batch 1, green for EBOVAC batch 2, blue for EBOPLUS, purple for VEBCON). Pearson correlation coefficients and p-values are shown. **B**. Genomic location of *LEF1-AS1* and *LEF1* on chromosome 4, showing antisense overlap and proximity to H3K27Ac histone marks and ENCODE candidate cis-regulatory elements (cCREs). **C**. Counts of *LEF1* and *LEF1-AS1* transcripts in the EBOLA vaccination. Cohorts shown as colors: red for EBOVAC batch 1, green for EBOVAC batch 2, blue for EBOPLUS, purple for VEBCON. On the x-axis are days post-vaccination, and on the y-axis are counts of *LEF1* and *LEF1-AS1*. **D**. Expression of *LEF1-AS1* and *LEF1* in response to other vaccines in independent datasets (GSE186006 and GSE196793 for influenza; GSE146658 for dengue) are shown as boxplots. On the x-axis are days post-vaccination, and on the y-axis are counts of *LEF1* and *LEF1-AS1*.

In summary, our multi-cohort analysis of transcriptomic responses to the rVSV-ZEBOV Ebola vaccine reveals a conserved and coordinated induction of long non-coding RNAs (lncRNAs) on day 1 post-vaccination. Among these, 11 lncRNAs showed consistent differential expression across cohorts and were tightly associated with immune gene expression signatures. Several lncRNAs, including *FAM225A, DOCK8-AS1*, and *LEF1-AS1*, demonstrated correlations with key regulators of the innate and adaptive immune responses, as well as with the magnitude and durability of vaccine-induced IgG titers. Functional annotation and genomic localization suggest that some of these lncRNAs, particularly *DOCK8-AS1* and *LEF1-AS1*, may act as promoter-antisense RNAs (PAS-RNAs), potentially influencing the expression of adjacent immune genes critical for T cell function and germinal center responses.

## Discussion

Vaccination triggers a tightly orchestrated sequence of immune events that begins immediately after antigen exposure. In the case of virus-based vaccines such as rVSVΔG-ZEBOV-GP, the response is typically characterized by a strong type I interferon (IFN-I) signature, largely driven by potent activation of myeloid cells (Vianello et al., 2022; Gonzalez Dias Carvalho et al., 2023). As a replicating viral vector vaccine, rVSVΔG-ZEBOV-GP may elicit a distinct early innate immune response compared to non-replicating viral vectors or inactivated virus vaccines. This early innate immune response includes TLR and NLR signaling in antigen-presenting cells, resulting in the expression of co-stimulatory molecules and cytokines that guide adaptive responses. Dendritic cells migrate to the lymph nodes, presenting antigens to naïve B and T cells. If the immune environment is sufficiently activated, naïve cells are retained in lymph nodes to scan for cognate antigens, initiating germinal center (GC) reactions. This process is resource-intensive and tightly regulated, involving T follicular helper (Tfh) cells that support B cell maturation, somatic hypermutation, and class switching from IgM to IgG (REF). The immune setpoint at the time of vaccination, can strongly influence the magnitude and quality of subsequent immune responses, with some individuals exhibiting a ‘naturally adjuvanted’ profile that mirrors early adjuvant-induced changes (Ramos, 2025). Our group has shown that rVSVΔG-ZEBOV-GP induces a sharp immune activation already on day 1 post-vaccination, likely driven by neutrophil clearance, dendritic cell migration, and naïve lymphocyte sequestration in lymph nodes; contrasting with the relatively quiescent immune landscape at day 0, which is necessary to ensure effective priming. ((Vianello et al., 2022; Gonzalez Dias Carvalho et al., 2023).While the dynamics of protein-coding genes in vaccine responses have been extensively characterized, less is known about the role of long noncoding RNAs (lncRNAs). A recent study (de Lima et al., 2019) demonstrated that vaccination elicits robust changes in lncRNA expression, identifying several noncoding transcripts that are tightly regulated post-immunization and that play active roles in modulating immune signaling pathways, including antigen presentation and B cell activation, highlighting lncRNAs as integral participants in shaping adaptive responses. In our study, we identified a conserved signature of 11 lncRNAs differentially expressed one day after rVSV-ZEBOV vaccination across three human cohorts and three of these (DOCK8-AS1, FAM225A, and LEF1-AS1) also correlated with patient’s antibody titer response. Among the other eigth lncRNA, *CYTOR* (Cytoskeleton Regulator RNA) has been linked to immunotherapeutic responses in cutaneous melanoma (Zhang et al., 2024), regulation of HIV infection (Kuzmina et al., 2024), and B-cell activation upon Epstein-Barr virus infection (Wang et al., 2019b) *DLEU2* (Deleted in Lymphocytic Leukemia 2) encodes a peptide that facilitates T-regulatory cell induction (Tang et al., 2024) and is associated with immune cell infiltration in hepatocellular carcinoma (Fu et al., 2021) and osteosarcoma (Fan et al., 2021). *FOXN3-AS1* (FOXN3 Antisense RNA 1) has been implicated in modulating tumor immune infiltration (Geng et al., 2022) and responses to immunotherapeutic treatment in acute myeloid leukemia (Ge et al., 2024), while *LINC00189* (Long Intergenic Non-Protein Coding RNA 189) shows correlated expression with CXCL2 in childhood obesity (He et al., 2024), and *LINC00487* (Long Intergenic Non-Protein Coding RNA 487) is associated with immune responses in B-cell lymphoma (Wang et al., 2022b). The remaining lncRNAs: *C8orf31* also known as LYS6-AS1, LY6S Antisense RNA 1) (Lyu et al., 2023), *SERPINB9P1* (Serpin Family B Member 9 Pseudogene 1)(Huang et al., 2022; Lv et al., 2025), and *ATP6V0E2-AS1* (ATP6V0E2 Antisense RNA 1(Kim et al., 2022a); have been reported in contexts such as cancer, stroke, and other pathological processes but lack direct immune-related characterization.

*DOCK8-AS1* emerged as a potential promoter-antisense RNA (PAS-RNA) regulating the expression of *DOCK8*, a gene essential for proper Tfh localization and B cell help in germinal centers (Deobagkar-Lele et al., 2024). *DOCK8* is an essential gene for correct Tfh cell positioning in the germinal center of lymph nodes in response to immune stimuli(Janssen et al., 2020). Mutations in DOCK8 are related to a severe immunodeficiency syndrome characterized by hyper-IgE production and recurrent viral and bacterial infections (Szczawinska-Poplonyk et al., 2011), as well as low antibody production and B cell maturation deficiency (aan de Kerk et al., 2013). We observed strong co-expression between *DOCK8-AS1* and *DOCK8* in both vaccinated volunteers and GRO-seq datasets of virus-infected lymphoblasts. Given that *DOCK8* deficiency is associated with impaired antibody class switching and susceptibility to infections (Biggs et al., 2017), the correlation between *DOCK8-AS1* levels and reduced IgG titers at multiple time points suggests a complex, possibly biphasic role in regulating humoral immunity.

While *LEF1-AS1* overexpression in patient blood has been implicated in cancer (Zheng et al., 2021) and COVID-19 pathogenesis (Vausort et al., 2025) it is often through a competing endogenous RNA (ceRNA) mechanism. It is known to act sponging miRNAs such as miR-489-3p (Zhang et al., 2023), miR-544a (Wang et al., 2019a), and miR-1285-3p (Zhang and Ruan, 2020). *LEF1-AS1*, was consistently downregulated on day 1 post-vaccination as it was its sense gene, *LEF1. LEF1* is crucial for lymphocyte development (Reya et al., 2000) and its regulation by *LEF1-AS1* may represent a transcriptional checkpoint to transiently suppress developmental programs and enable antigen-driven B cell activation. This downregulation, according to our data, was unique to Ebola vaccination and was not observed in other vaccine datasets (influenza, dengue), which could point to a potentially vaccine-specific regulatory mechanism.

The lncRNA *FAM225A* was positively correlated with IgG titers on days 14 and 56 post-vaccination. It also showed strong correlation with interferon-stimulated genes such as IRF7, IFIH1, and USP18, the latter of which is required for VSV-ZEBOV efficacy in macrophage-mediated response (Pinski and Messaoudi, 2021). These data suggest that *FAM225A* may contribute to the initiation of an effective antiviral response and facilitate the transition to a durable adaptive immunity via Tfh and GC support.

Collectively, our findings reveal that lncRNAs play a significant and coordinated role in shaping the immune landscape following rVSV-ZEBOV vaccination. They appear to modulate both early innate signaling and the quality of the subsequent antibody response, acting through mechanisms such as cis-regulation of immune effectors and PAS-RNA-mediated transcriptional control. This is the first study to highlight the role of lncRNAs in the quality and longevity of the humoral response to Ebola vaccination, and it opens the door for future mechanistic investigations and vaccine-enhancing interventions that target noncoding RNA networks.

## Methods

### Volunteer’s cohorts

Volunteers from the VSV-Ebovac and VSV-EBOPLUS cohorts were participants of the clinical trials for the rVSVΔG-ZEBOV-GP Ebola vaccine conducted in Europe and North America. VSV-Ebovac: phase 1/2, randomized, double-blind, placebo-controlled, dose-finding trial in Geneva, Switzerland (November 2014, to January 2015; NCT02287480). VSV-EBOPLUS: phase 1b, randomized, double-blind, placebo-controlled, dose-response trial in the USA (Dec 5, 2014, to June 23, 2015; NCT02314923). The VEBCON volunteers were participants of the rVSV-ZEBOC clinical trial conducted in Europe: a phase 1, open-label, dose-escalation single-center trial conducted in Hamburg, Germany (Nov 2014 to Dec 2014).

### Blood samples and RNA extraction and RNA sequencing

For the VSV-Ebovac and VSV-EBOPLUS cohorts, 2.5 mL venous blood was collected in PAXgene blood RNA tubes (PreAnalytiX, Hombrechtikon, Switzerland) from all participants on days 0, 1, 2, 3, and 7. Comprehensive methods for the extraction and sequencing have been described before (Vianello et al., 2022).

### Public data obtention and experimental methods

For the Vebcon cohort, raw RNA sequencing counts and patient metadata were obtained on the Gene Expression Omnibus (GEO) database (GEO accession: GSE97590) (Rechtien et al., 2017). Briefly, RNA was extracted from the blood of participants collected on days 0, 1, 3, and 7 after vaccination. RNA-seq libraries were generated using the NEBNext Ultra RNA Library Prep Kit for Illumina (NEB) and sequenced on the Illumina HiSeq 2500 (single read, 50-bp run). Reads were aligned to the human reference assembly (GRCh38.p7) using STAR (v2.4.2a) and quantified with “–quantMode GeneCounts”. Gene annotation was obtained from Ensembl (release 79, http://www.ensembl.org).

### IgG quantification

ELISA assays to detect ZEBOV-GP-specific IgG antibody titers were performed for previously published studies (Regules et al., 2017). Briefly, ZEBOV glycoprotein-specific antibodies were quantified with the Filovirus Animal Non-Clinical Group (FANG)-approved ELISA by use of the homologous Zaire-Kikwit strain glycoprotein, following USAMRRIID’s standard operating procedure (SOP AP-03-35; USAMRIID ELISA).

### Differential expression (DE) analysis

Genes with less than 1 count in 10 or more samples were removed from the raw count tables prior to DE analysis. The R package *edgeR* was used to perform DE using the *glmQLFit* function after using the *calcNormFactors* function to normalize counts and the *estimateDisp* function to estimate dispersions (Chen et al., 2025). DE genes were considered those with adjusted p-value lower than 0.05 and absolute log2FoldChange higher than 0.26 (absolute fold change of 20%). DE testing was performed between the samples from volunteers on days 1, 2, 3, or 7 against their baseline levels on day 0 (prior tobefore vaccination). The R package *biomaRt* was used to convert gene name synonyms from the VSV-EBOVAC and VSV-EBOPLUS DE results to HUGO Gene Nomenclature Committee (HGNC) format. VEBCON DE genes were converted from Ensembl gene ID to HGNC format. The GENCODE v37(Frankish et al., 2021)) human genome annotation was used to obtain gene type labels for each HGCN ID. Only genes labeled as “lncRNA” according to the GENCODE annotation were kept for further DE analyses. DE analysis results were plotted using the R package *ggplot2* and tidyomics (Hutchison et al., 2024).

### Correlation meta-analysis

Baseline normalized (Basenorm) expression values were calculated for each day after the vaccine for all volunteers in all cohorts. Basenorm consisted of subtracting day 0 expression values from the expression values on each of the subsequent days following the vaccine for each patient. Basenorm expression values of volunteers in each cohort were used to perform pairwise Pearson’s correlation tests (*cor*.*test* function in R) between selected lncRNAs and all other genes available in the Basenorm expression matrices. Basenorm values were also used to perform pairwise Pearson’s correlation tests (*cor*.*test* function in R) between selected lncRNAs and IgG titers on all measured days after vaccination for the VSV-EBOVAC and VSV-EBOPLUS cohorts. After, the R package *metafor* (Viechtbauer, 2010)was used to perform a random-effects model meta-analysis of lncRNA-gene and lncRNA-IgG correlation p-values and correlation coefficients across all cohorts. Correlation pairs between lncRNAs and genes with a meta-adjusted p-value lower than 0.05 were considered significant. Correlation pairs between lncRNAs and IgG titers with a meta-p value lower than 0.05 were considered significant.

### Functional enrichment analysis

Genes with a significant meta-p value correlation were used to perform overrepresentation analysis (ORA) using the Reactome pathways (Gillespie et al., 2022) and Blood Transcription Modules (BTMs) (Altman et al., 2021) databases as reference. For the correlation ORA, the genes were split into positively or negatively correlated for each lncRNA according to the meta-correlation coefficient values. Targets of miRNAs were submitted to separate ORAs for each miRNA. The function *fora* of the R package *fgsea* was used to perform ORA. All genes in the filtered expression tables for each cohort were used as the background universe for the ORA. Significantly enriched pathways were those with an adjusted-p -value lower than 0.05. All correlation and enrichment analysis results were plotted using the R packages *ggplot2* and *pheatmap*.

### Global run-on sequencing (GRO-seq) analysis

Raw GRO-seq reads were downloaded from the Sequence Reads Archive (SRA) (Project accession: PRJEB22484). Briefly, human GM12878 lymphoblasts were infected *in vitro* with Sendai Virus Cantell Strain at 3000 HA units per 1 million cells (or at ∼1×10e8 EID50). To obtain nascent RNAs for sequencing, a previously published GRO-seq protocol was used (Bitzer et al., 1999). Nuclear run-on (NRO) reaction was performed with isolated nuclei treated with BrU for labeling of nascent RNAs. BrU incorporated nascent RNA from total nuclear RNA was purified using Br-UTP beads. The RNA was reverse transcribed into cDNA and sequenced on the Illumina Genome Analyzer II according to the manufacturer’s instructions with a small RNA sequencing primer. Detailed descriptions of the methods used are available at SRA accession **ERX2171558**. Computation GRO-seq analysis was performed according to the guidelines of a previously published methods review (Nagari et al., 2017). Raw GRO-seq reads were trimmed to remove adapters and poly-A tails using Cutadapt and aligned to the human genome (GRCh38.p13) using the Burrows-Wheeler Aligner (BWA). Sorted .bam files were generated from the BWA output .sam files using Samtools. The .bam files were filtered using Samtools to extract the genomic region of DOCK8-AS1 and DOCK8, before downstream analysis. The R package *Rsamtools* was used to import filtered .bam files to R, and an in-house script was used to extract and plot GRO-seq counts on the genomic region of the DOCK8-AS1 and DOCK8 genes. The R package *ggbio* was used to plot gene transcript structures.

### Systematic search for other vaccination datasets

To assess whether the transcriptional regulation of *LEF1-AS1* and its association with *LEF1* expression were specific to the Ebola vaccine, a systematic search of publicly available transcriptomic datasets from other human vaccination studies was conducted searching for RNA-seq studies that included peripheral blood samples collected at multiple time points following vaccination with other viral vaccines in Gene Expression Omnibus (GEO). Selected studies had public gene-level expression matrices and sample metadata, and multiple timepoints. No study with a day 1 time point was found. Four relevant datasets were found: **GSE146658** (dengue vaccine), **GSE274606, GSE196793**, and **GSE186006** (influenza vaccines). GSE274606 had no *LEF1-AS1* expression, so it was excluded. Metadata containing vaccine time points or sample identifiers was integrated into the expression tables. All visualizations were generated using ggplot2 and tidyomics (Hutchison et al., 2024).

